# Remating responses are consistent with male post-copulatory manipulation but not reinforcement in *D. pseudoobscura*

**DOI:** 10.1101/072066

**Authors:** Jeremy S. Davis, Dean M. Castillo, Leonie C. Moyle

## Abstract

Reinforcement occurs when hybridization between closely related lineages produces low fitness offspring, prompting selection for elevated reproductive isolation specifically in areas of sympatry. Both pre-mating and post-mating prezygotic behaviors have been shown to be the target of reinforcing selection, but it remains unclear whether remating behaviors experience reinforcement, although they can also influence offspring identity and limit formation of hybrids. Here we evaluated evidence for reinforcing selection on remating behaviors in *D. pseudoobscura*, by comparing remating traits in females from populations historically allopatric and sympatric with *D. persimilis*. We found that the propensity to remate was not higher in sympatric females, compared to allopatric females, regardless of whether the first mated male was heterospecific or conspecific. Moreover, remating behavior did not contribute to interspecific reproductive isolation among any population; that is, females showed no higher propensity to remate following a heterospecific first mating than they were following a conspecific first mating. Instead, we found that females are less likely to remate after initial matings with unfamiliar males, regardless of species identity. This is consistent with one scenario of postmating sexual conflict in which females are poorly defended against post-copulatory manipulation by males with whom they have not co-evolved. Our results are generally inconsistent with reinforcement on remating traits, and suggest that this behavior might be more strongly shaped by the consequences of local antagonistic male-female interactions than interactions with heterospecifics.

## Introduction

Because hybridization between incompletely isolated species can be costly—in terms of reduced fecundity or offspring survival or fertility—selection is expected to favor traits that reduce the frequency or consequences of these matings in nature (Dobzhansky 1940). This ‘reinforcement’ of incomplete reproductive isolation is thought to play a key role in speciation, especially where there is secondary contact between close relatives (Ortiz-Barrientos et al. 2009). Reinforcement has frequently been examined in the context of selection on premating traits, such as courtship displays or behaviors, which can act to prevent heterospecific matings (e.g., Saetre et al. 1997, Rundle and Schluter 1998). However, post-mating traits could also be subject to reinforcing selection (Servedio and Noor 2003) as could traits that integrate pre- and post-mating responses, such as postmating control of paternity via variable remating rate (Marshall et al. 2002, Kisdi 2003). Control over mate and paternity choice has been shown to evolve rapidly in response to antagonistic coevolution between the sexes (e.g., Rice 1996, Miller and Pitnick 2002, Manier et al. 2013a). Such rapidly evolving reproductive traits can potentially drive divergence between populations and might contribute strongly to reproductive isolation (Parker and Partridge 1998, Rice 1998, Howard 1999, Gavrilets 2000, Panhuis et al. 2001, Martin and Hosken 2003, Ritchie 2007, Howard et al. 2009, Manier et al. 2013b). Therefore both mating and remating behaviors are potentially interesting candidates for examining the evolution of isolating mechanisms between species, including in the context of reinforcement.

One key expectation under reinforcement is that populations that are historically sympatric with closely related heterospecifics will show stronger isolation than populations that are historically allopatric (Butlin 1987, Servedio and Noor 2003). This is because only sympatric populations will have experienced selection to avoid producing lower fitness hybrid offspring. Mate choice during the first mating has been observed to show patterns consistent with reinforcement (e.g., Noor 1995, Higgie et al. 2000, Saetre et al. 1997, Rundle and Schluter 1998), whereby sympatric females are more discriminating against heterospecifics than are allopatric females. In comparison to initial mate choice, whether remating rates respond to reinforcement is largely unknown (Marshall et al. 2002; but see Matute 2010, and Discussion). Decreasing the time to remating (latency) or increasing the propensity to remate allows females to manipulate paternity, including after mating with a suboptimal male (variously called a ‘rescue effect’ (Fricke et al. 2006) or the ‘trading up’ hypothesis (Byrne and Rice 2005)). Because mating with heterospecifics is generally suboptimal, remating rate could respond to reinforcing selection such that sympatric females increase their propensity to remate with conspecifics following a heterospecific mating (Marshall et al. 2002). It is also possible that exposure to heterospecifics could generally increase remating rates of females in such populations, regardless of first male identity. In comparison, females from populations that are geographically allopatric are not expected to elevate remating responses.

Nonetheless, making predictions about remating rate is complex because remating behaviors are the product of both female choice and male manipulation. For example, in *Drosophila,* females are known to exhibit cryptic female choice by controlling number of mates and/or by preferentially using sperm from some male partners (Manier et al. 2010, Lupold et al. 2013, Manier et al. 2013c). In turn, male *Drosophila* seminal fluid proteins transferred during copulation are known to suppress female remating rate, increase oviposition rate, and reduce lifespan, potentially resulting in net fitness reductions for females (Parker and Partridge 1998, Sirot et al. 2009, and references therein). The resulting antagonistic male-female coevolution acting on these traits can lead females to be poorly defended against males with whom they have not co-evolved (Rice 1998, Parker and Partridge 1998). Under this scenario, for example, allopatric females that are less equipped to defend against heterospecific encounters might exhibit reduced remating rates, even when remating would be individually beneficial. It can, however, be difficult to make general predictions about the direction of female responses to unfamiliar mates, because this is expected to depend on which sex is “ahead” in the coevolutionary arms race, which can vary depending upon the precise details of these male-female interactions (Long et al. 2006, reviewed Tregenza et al. 2006, and see Discussion).

We sought to examine whether remating rates might respond to reinforcing selection in a *Drosophila* species pair that is a canonical example of reinforcement of premating isolation. *Drosophila persimilis* and *Drosophila pseudoobscura* are recently diverged (500 kya) sister species with distinct but significantly overlapping ranges (Shaeffer and Miller 1991, Wang et al. 1997, Machado et al. 2002). *D. pseudoobscura* has a wide geographic range in North America, stretching west from the Pacific to close to the Mississippi River and far South into Central America; *D. persimilis* has a far narrower range completely sympatric with *D. pseudoobscura* and not extending farther east than the Sierra Nevada and Cascade Mountain ranges (Figure 1). These species exhibit incomplete reproductive isolation and hybridize in the laboratory; natural hybrids, while rare, have been found in the wild (Dobzhansky 1973, Kulathinal et al. 2009). In addition, mate choice patterns consistent with reinforcement have been directly demonstrated in this species pair, whereby allopatric *D. pseudoobscura* females mate at a higher rate with *D. persimilis* males than do *D. pseudoobscura* females from sympatric populations (Noor 1995), although see Anderson and Kim (2005, 2006) for more complex patterns of isolation between sympatric and allopatric populations. Additionally, a recent study evaluating other components of reproductive isolation in this species pair (Castillo and Moyle 2016) found no difference in first mating rates between allopatric and sympatric *D. pseudoobscura* paired with *D. persimilis* males. The well-established ranges and prior focus on evaluating reinforcement in this species pair make it particularly suited for examining whether remating rate might also respond to reinforcing selection.

**Figure 1:**
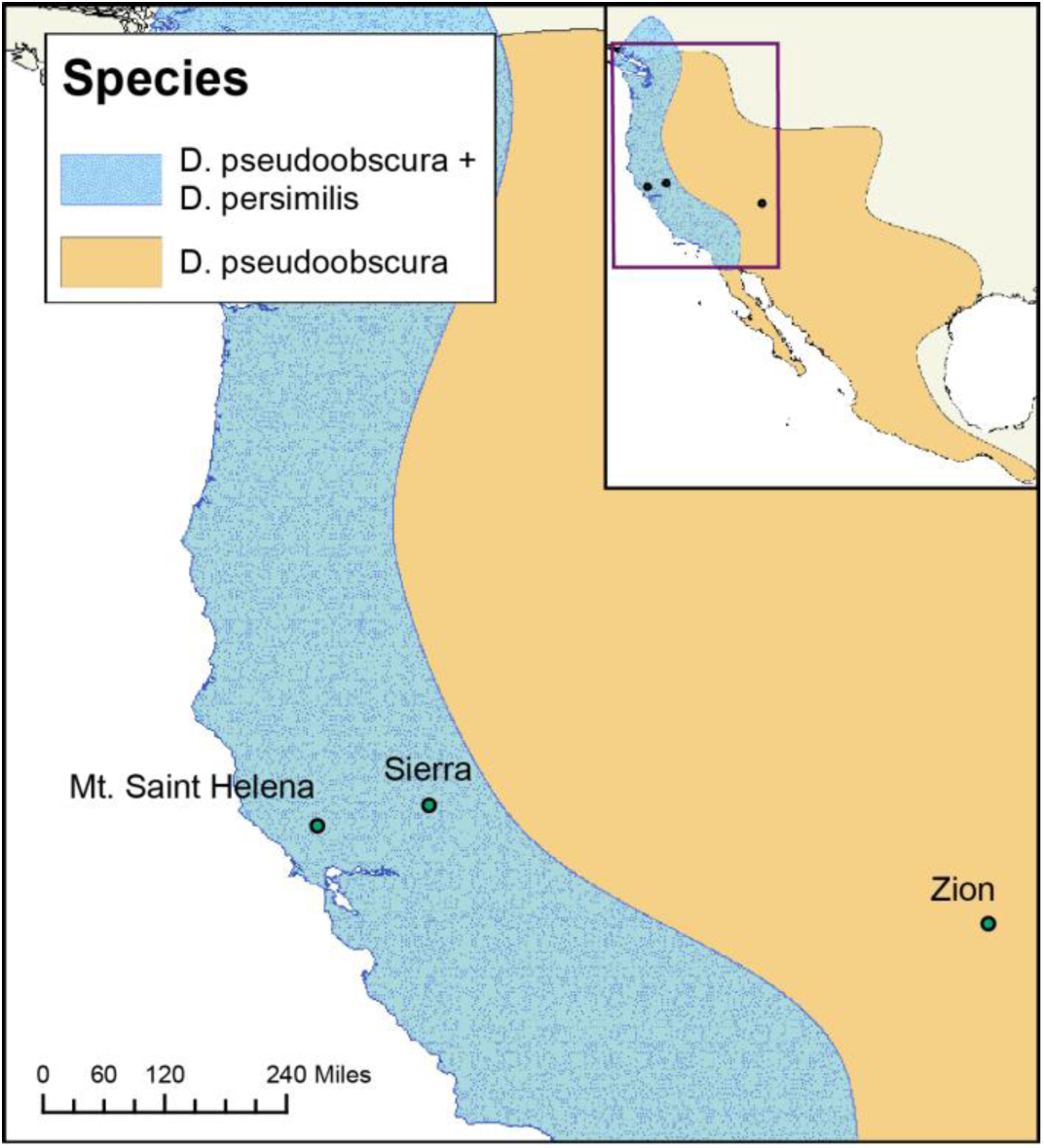
Collection locations for *Drosophila pseudoobscura* and *D. persimilis* study populations. Mt. Saint Helena and Sierra are sympatric locations (both species); Zion is an allopatric site (*D. pseudoobscura* only). Inset: North American range maps for the two species; the range of *D. persimilis* is entirely contained within the broader *D. pseudoobscura* range.

Our primary goal in this study was to evaluate evidence for reinforcing selection on remating behaviors of *D. pseudoobscura* females, using populations historically allopatric and sympatric with *D. persimilis*. To do so, we evaluated mating traits in females from three target *D. pseudoobscura* populations: two populations sympatric with *D. persimilis*, and one allopatric population (Figure 1).

These populations were a subset of those examined in a larger parallel study of patterns of first mating and conspecific sperm precedence between these species (Castillo and Moyle 2016). Following a first mating with either a heterospecific or a conspecific male, females were given the opportunity to remate with a male from their own population. We assessed whether female remating response depends on identity of the first mated male and, specifically, whether the propensity to remate depends upon female population identity (allopatric/sympatric). If remating behaviors have evolved in response to the presence of heterospecifics, we expected that *D. pseudoobscura* females from sympatric sites would more readily remate if their first mating was with a heterospecific male, consistent with an evolved response to limit the number of hybrid offspring sired from this first mating. An alternative expectation is that females from sympatric populations remate at a higher rate irrespective of first male identity, as a simpler response to potentially suboptimal first matings.

## Methods

### D. pseudoobscura *and* D. persimilis *collection and maintenance*

All stocks were reared on standard media prepared by the Bloomington Drosophila Stock Center, and were kept at room temperature (~22C). We used a subset of isofemale lines from a larger panel that were collected in the summers of 2013 and 2014 at three sites (Figure 1). Allopatric *D. pseudoobscura* were collected at Zion National Park, UT (kindly provided by N. Phadnis). Sympatric *D. pseudoobscura* and *D. persimilis* were collected at two sites: Mt. St. Helena, CA (*D. pseudoobscura* collected by A. Hish/M. Noor, and *D. persimilis* collected by D. Castillo); and, near Meadow Vista and Forest Hill, CA (called here ‘Sierra’; Figure 1) (*D. pseudoobscura* and *D. persimilis* collected by D. Castillo). For both sympatric populations, both species were present in field collections and can be considered truly co-occurring/sympatric. Our three focal populations are a subset of four populations used in a parallel study that evaluated evidence for reinforcement on first matings, and on conspecific sperm precedence (Castillo and Moyle 2016). (The current study excludes analysis of an additional allopatric population from Lamoille Canyon, NV). All but one of the 6 isofemale lines from our three populations are shared in common with the other study (MSH3 is not used in Castillo and Moyle 2016), enabling us to compare remating data from both experiments here (see below), as well as reassess the prior first mating result with data obtained from our first mating observations.

### Mating and Remating assay

To examine remating behaviors in females from our three target *D. pseudoobscura* populations, we used a design in which each female was initially paired with 1 of 5 different types of male (males from each of the 3 *D. pseudoobscura* populations and 2 *D. persimilis* populations). Five day-old virgin females were transferred individually without anesthesia to vials with individual 5-day old virgin males and allowed to mate for 24 hours before the male was removed. Females were then allowed to lay for 9 days, a refractory period that pilot trials indicated gives ample time for females to become receptive to males again. Those that produced larvae (and therefore were guaranteed to have mated with the first male) were then given the opportunity to remate with a second, 5-day-old virgin male. The second male was always from the same population as the target female, to ensure females would mate most readily during the second mating. This procedure was performed for each combination of our three female populations and five first male types (15 total cross combinations). Two complete experimental blocks were performed for each cross combination, using two unique isofemale lines from each population.

Within each experimental block, a minimum of 8 biological replicates were carried out for each combination of first and second male matings.

For each first male pairing, mating behavior was directly observed for 3 hours, and copulation latency (time to start of copulation) and duration (time from start to end of copulation) were recorded. Following the 3-hour observation period, pairs were maintained together for an additional 21 hours, and vials were checked 7 days later for larvae to determine if mating occurred within first 24 hours but outside the initial 3-hour observation window. This allowed us to assess whether female population origin influences mating behavior in the first male mating, and whether this varied according to male population identity. For each second male pairing, female mating behavior was assessed in terms of copulation latency and mating duration within the first 3 hours of pairing. This allowed us to evaluate whether females vary their remating behavior in response to the population and/or species identity of their first mate, in addition to whether these responses differed in females from allopatric versus sympatric sites. Finally, differences among isofemale lines in overall propensity to remate following conspecific first matings, was used to confirm that there was heritable genetic variation for this trait within *D. pseudoobscura* (Results). Detailed mating procedures are provided in Supplementary material.

After completing at least 8 replicates, we found that copulation duration during the first mating was indistinguishable among all crosses, and copulation latency was either similarly rapid (<10 minutes) in all conspecific pairings, or inconsistently and rarely observed within the first 3 hours in heterospecific pairings (Results). Based on these findings, for the remaining 14 replicates (which primarily focused on heterospecific second pairings) first matings were no longer directly observed for the first 3 hour period, but were instead simply scored for presence/absence of larvae 7 days after co-housing each male-female pair for 24 hours. Regardless of this change for first matings, remating behavior was always assessed as observed copulation, and copulation latency and duration, within the first 3 hours of co-housing.

To assess whether detected remating differences could be explained by differences in sperm usage and depletion between different cross types, we tracked progeny production of 2 isofemale lines, one allopatric and one sympatric *D. pseudoobscura* strain, across 7 days. As with all first matings, individual virgin females were mated overnight with either a male from their own population, a *D. pseudoobscura* male from a different population, or a *D. persimilis* male. Males were removed after 24 hours. We found no significant differences in number of progeny produced from own population males, conspecific males from a different population, or heterospecific males, for either allopatric or sympatric isofemale lines, consistent with a previous study that found no evidence for non-competitive gamete isolation contributing to reproductive isolation in this species pair (Lorch and Servedio 2005). Allopatric females produced an average of 86 progeny, which did not differ based on whether they mated with males from their own population versus different population conspecifics (β = 9.400; P = 0.603) or versus heterospecifics (β = 7.487; P = 0.609). Similarly, there was no difference in the number of progeny produced for sympatric females (77 progeny) when mated with males from their own population versus different population conspecifics (β =2.667; P = 0.913) or versus heterospecifics (β =7.667; P = 0.702)

Although Castillo and Moyle (2016) did not observe remating directly, data from that experiment can be used to glean some additional information about remating rates in sympatric versus allopatric females. Similar to the design here, in that study virgin *D. pseudoobscura* females were housed with *D. persimilis* males for 24 hours and then, following a period of 7 days, were given the opportunity to remate with *D. pseuodoobscura* males. Progeny after this second mating were scored (*D. persimilis* male was marked with a visible marker, and hybrid males are sterile), providing information on whether females remated or not. Females are inferred to have failed to remate if all progeny after second mating were hybrid; that is, if all males were sterile and all females carried the visible mutation). These data were used as an additional test of whether allopatric and sympatric females differed in their propensity to remate (see results).

### Statistics

A χ2 test of independence was used to compare overall *D. pseudoobscura* female mating rates in first pairings with conspecific versus heterospecific males. To make more specific comparisons among groups, we used logistic regression on presence/absence of larvae after mating (mating was considered a binary variable). Logistic regressions were used to assess differences in the mating probabilities of all females during their first matings, and during remating trials, depending upon whether they were initially paired with first males of three classes: males from their own population, males from a different conspecific population, or heterospecific males. Probabilities of mating and of remating were also specifically compared between *D. pseudoobscura* females historically allopatric and sympatric with *D. persimilis*. For all logistic regressions, differences between mating types were inferred by examining significance of the regression coefficients. Negative coefficients signified categories where matings were less likely to occur, and positive coefficients signified that mating was more likely to occur.

To analyze quantitative copulation latency, we primarily used Cox proportional hazard models in the *survival* package (Therneau 2013) in R, which let us take into account the mating and remating rates as well as probability of mating within our 3 hour observation. For one comparison (allopatric versus sympatric remating latency) we used parametric survival regression (see Supplement for details). We included female genotype in the proportional hazard models to account for correlated observations within a given isofemale line (see Supplemental information for details). Survival curves for a specific mating category were considered different when the coefficient from the model was significantly different than the zero. Negative coefficients signified categories where matings occurred more slowly than baseline, and positive coefficients signified that mating occurred more quickly than baseline. Baseline was always mating involving males from the females own population. Finally, a χ2 test of independence was used to compare overall *D. pseudoobscura* remating rates between allopatric and sympatric females, using the remating data integrated from Castillo and Moyle (2016).

## Results

### Initial mate choice contributes to reproductive isolation between species but is not stronger in sympatry

We confirmed that *D. pseudoobsura* females discriminate against *D. persimilis* males; while almost all conspecific matings were successful (164/168), only 25% of heterospecific pairings resulted in mating (93/369), a significant difference in mating propensity (χ2 = 239.70; *P* < 2.2×10^−16^). Logistic regressions similarly indicated that the proportion of heterospecific matings was significantly lower than the proportion of first matings either with males from a different conspecific population (β = 3.5927; *P* = 8.31×10^−10^) or with males from the females own population (β = 4.0073; *P* = 7.13×10^−05^) (Figure 2). There was no difference in the propensity to mate of females paired with males from their own population versus males from different conspecific populations (β = 0.4146; *P* = 0.722) (Figure 2), indicating that female choice in conspecific first matings was not sensitive to the population origin of the conspecific male.

**Figure 2:**
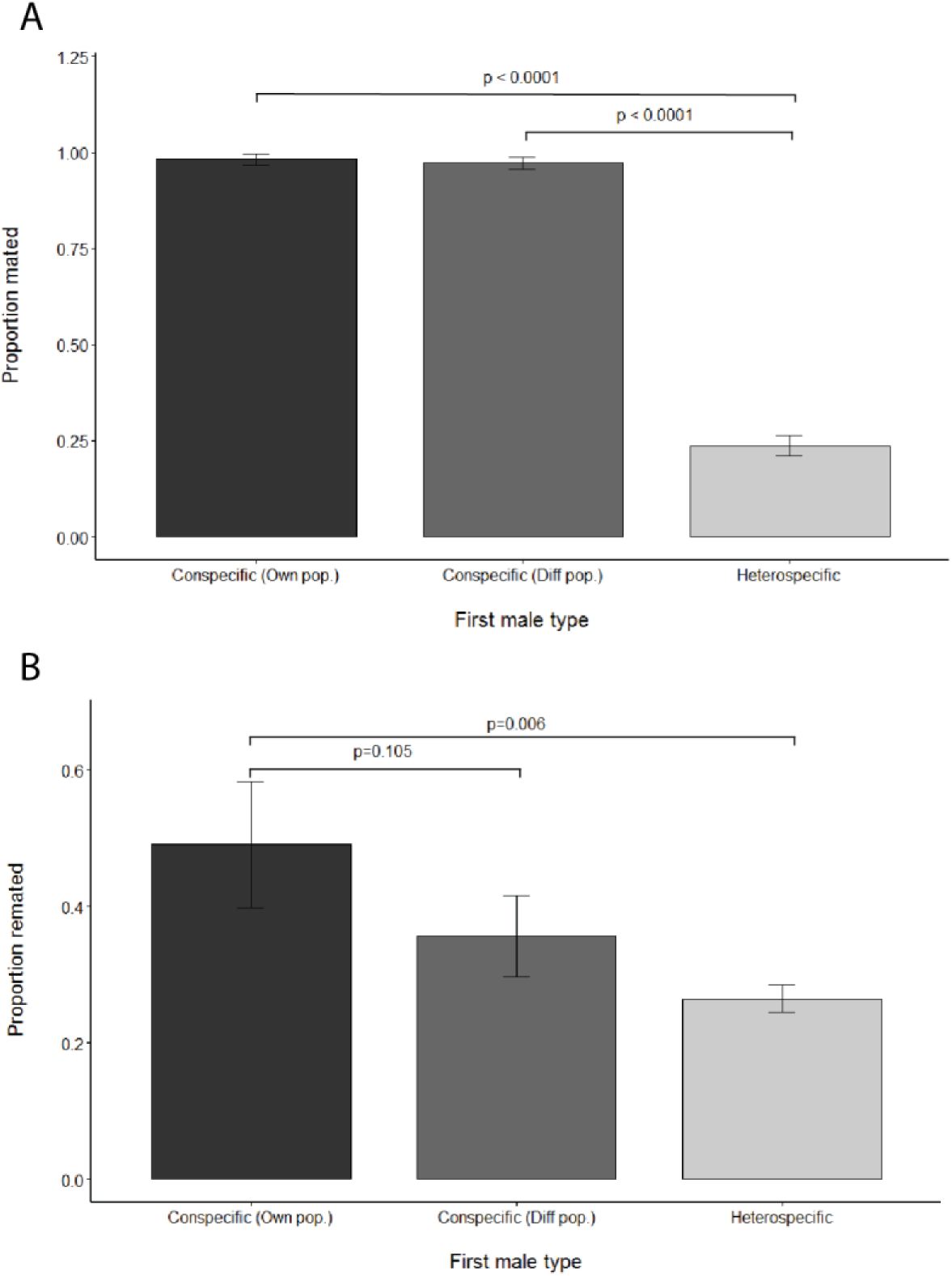
Mating (panel A) and remating (panel B) probabilities of *D. pseudoobscura* females following first matings with males from their own population (Own pop), a different conspecific population (Diff pop), and heterospecific males. P-values are from logistic regressions (see Results).

**Figure 3:**
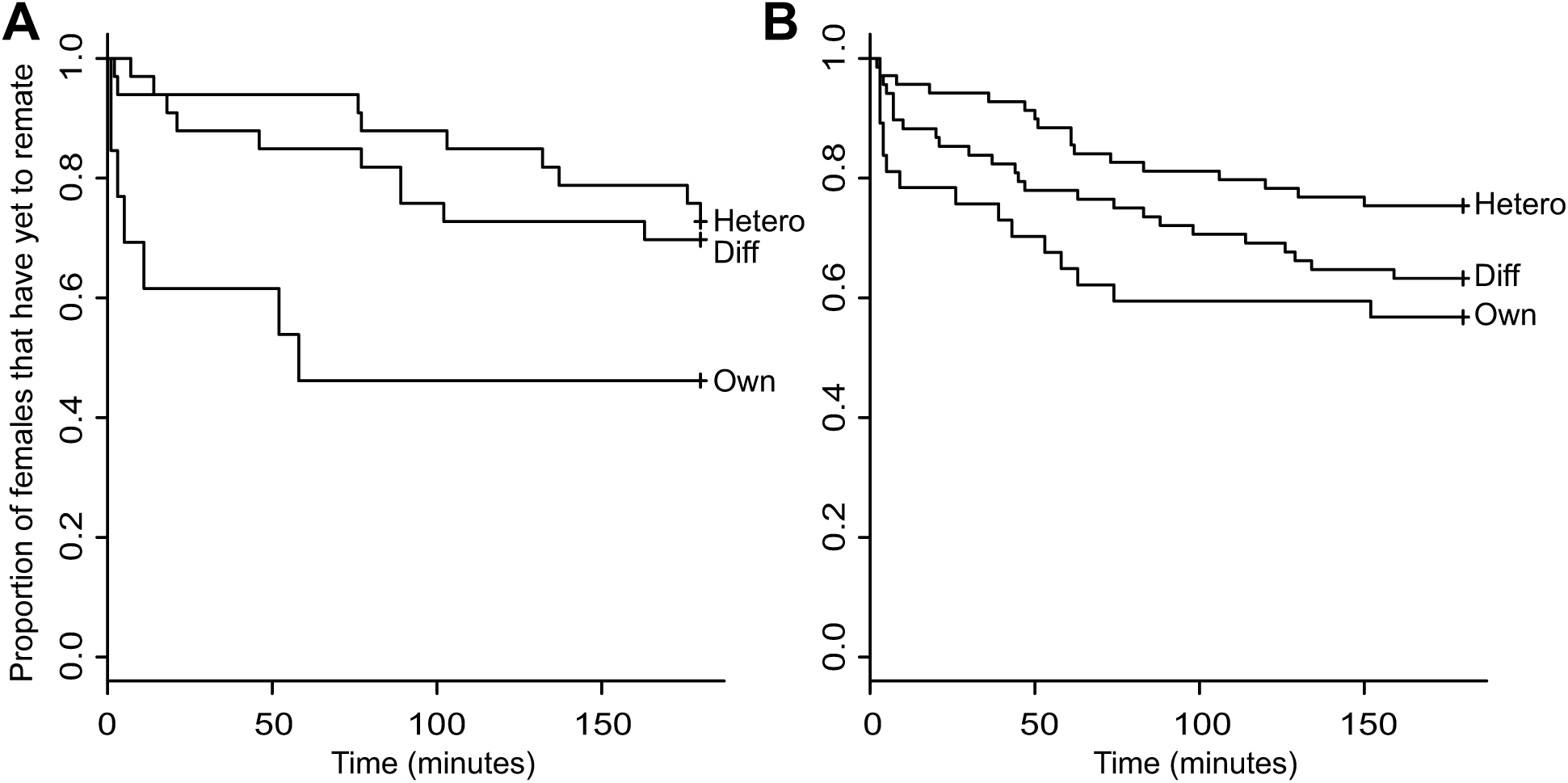
Survival curves showing remating latencies of allopatric (A) and sympatric female (B) *D. pseudoobscura* when mated to different classes of first males: Own (conspecific male from the same population as the female); Diff (conspecific male from a different population); Hetero (heterospecific male). Figure S1 shows survival curves of remating latency separately for each isofemale line.

To test for patterns consistent with reinforcement on first mating, we fit a logistic regression to first mating success according to whether female *D. pseudoobscura* were from a population that was allopatric or sympatric with *D. persimilis*. We found no significant difference between allopatric and sympatric females in probability of mating with a heterospecific first male (β = −0.1079; *P* = 0.623), consistent with prior observations of mating patterns that used more isofemale lines and one additional allopatric population comparison (Castillo and Moyle 2016). We did not analyze differences in copulation latency between allopatric versus sympatric females in heterospecific first matings because too few of these mating events occurred within the directly observed first 3 hours of cohousing. Only 16 of 133 directly observed heterospecific pairings resulted in copulations within the first 3 hours, corresponding to 14% and 15% of sympatric and allopatric female pairings. An additional 20 of the 133 directly observed pairings resulted in progeny, but these matings occurred within the subsequent (unobserved) 21 hour period of co-housing.

### Sympatric females are slower to mate with conspecific males in first matings

If exposure to heterospecifics has resulted in general changes in intrinsic mating behavior, rather than specific responses directed at heterospecific genotypes, allopatric and sympatric females should differ in mating responses to conspecific males. Allopatric and sympatric females did not differ in probability of mating when paired with conspecific males (regardless of their population of origin) (β = - 17.22; *P* = 0.994). However, females from sympatric populations took significantly longer to initiate copulation with conspecific males than did allopatric females, despite persistent courtship by males (β = −0.2693; *P* = 0.0096). When we simultaneously tested for an effect of sympatry and for the population of origin of the conspecific male, mating latency did not differ according to the specific population of the conspecific male (β = 0.0584; *P* = 0.6717) but the difference between sympatry and allopatry remained (β = =-0.2720; *P* = 0.0076). In other words, sympatric females are slower to initiate copulation, regardless of the population identity of the first conspecific male (i.e. own versus other conspecific population) with which they are paired. It is possible that this is a subtle behavioral response to past selection imposed by heterospecifics: if female *D. pseudoobscura* in sympatry have adapted to encountering heterospecific males, they might be more circumspect in their initial mating decisions in general. This longer latency might contribute to fewer accidental heterospecific matings, especially under less restrictive conditions than those imposed by our lab co-housing experiment.

### Remating varies depending on the identity of the first mating male, but does not contribute to interspecific reproductive isolation or to enhanced isolation in sympatry

To evaluate whether *D. pseudoobscura* females differed in their readiness to remate depending on the identity of the first male they mated with, we compared the frequency of remating and the copulation latency in remating trials following three classes of first mating: with conspecific males from their own population, with conspecific males from a different population, or with heterospecific males. We found that analyses of both mating probability and latency to copulation indicate that remating happens more readily when females first mate with familiar (own population) males, than when initially mated with unfamiliar conspecifics or with heterospecifics. In terms of remating probability, females initially mated to their own population males were significantly more likely to remate compared with females initially mated to a *D. persimilis* male (β = −0.98291; *P* = 0.00555), although the probability of remating did not differ significantly between females previously mated with conspecific males from their own population versus from a different conspecific population (β = −0.55603; *P* = 0.10471). In terms of latency to copulation, females first mated with their own male remated more quickly (had shorter latency) than females initially mated with either conspecifics from different populations (β = −0.5213; *P* = 0.02195) or heterospecifics (β = −0.8035; *P* =0.000526). (Although trending in this direction, copulation latency was not significantly shorter in females initially mated with conspecifics from a different population versus with heterospecific males (i.e., the confidence intervals on β coefficients overlap).) These observations also indicate there is no generalized female *D. pseudoobscura* response to increase remating following heterospecific first matings.

To test for patterns consistent with reinforcement on remating, we assessed whether allopatric versus sympatric females differ in their remating behaviors following first matings with *D. persimilis* males. We found that they did not differ in their probability of remating (β = −0.2851; *P* = 0.5447), or in how rapidly they remated (parametric survival regression; β = 0.252; *P* = 0.5280), following a heterospecific first mating. Finally, using a second set of mating data from Castillo and Moyle (2016), we examined the number of females that failed to remate compared to the total number of remating trials scored, and found there was no significant difference in remating rate between females from allopatric versus sympatric populations (χ2=0.1445; df=1, *P*=0.7029). Note that we detected significant differences among *D. pseudoobscura* isofemale lines in their overall propensity to remate following a conspecific first mating (Wald’s χ2; df = 5; *P* = 0.0352), indicating there is genetic variance for remating behavior available to selection in this species.

### Allopatric and sympatric females do not differ in remating behavior with conspecifics

To investigate whether allopatric versus sympatric females differ in their intrinsic propensity to remate, we compared remating probability and latency between allopatric and sympatric females that had first mated with conspecifics; we found that they did not differ in their probability of remating (β = 0.1586, *P* = 0.6569) or in their latency to copulate in remating trials (β = 0.1616, *P* = 0.5840). When we simultaneously tested for an effect of sympatry and for the population of origin of the first mated conspecific male, allopatric and sympatric females still did not differ in remating latency (β = 0.0961, *P* = 0.7437); however, we did detect a first male population effect, such that remating occurred more rapidly when females had mated first with a conspecific from their own population (β = −0.4975; *P* = 0.0239). This is consistent with our findings that females overall mate quickest following own-male first matings (above). Sympatric and allopatric females did not differ in remating latency following own male matings (β = −0.3492, *P* = 0.1570).

## Discussion

In this study our primary goal was to evaluate if sympatric *D. pseudoobscura* females remate more quickly or at a higher rate when previously mated to a heterospecific *D. persimilis*, as expected if remating behavior has responded to reinforcing selection in sympatry. We found no evidence for reinforcement effects on remating, in either probability of remating or in latency to copulation, when females had previously mated to heterospecifics. Sympatric females were also no more likely or faster to remate after conspecific first matings. Therefore our results indicate little evidence that remating behavior in our sympatric populations has responded specifically to reinforcing selection. In addition, our results also imply that our sympatric *D. pseudoobscura* females do not show a generalized change in remating behavior (either an increased general propensity to remate or to remate more quickly) in order to minimize the consequences of suboptimal (especially heterospecific) matings. Our results differ from the only other study (of which we are aware) to compare remating rates between females allopatric and sympatric with a closely-related conspecific species. In it, Matute (2010) found that *D. yakuba* females sympatric with *D. santomea* exhibit greater remating rates after heterospecific matings, compared to *D. yakuba* females that are allopatric, a pattern that is consistent with the expectations of reinforcement on remating, but that could also be explained by less direct effects (see below).

Given that there is genetic variation for *D. pseudoobscura* female remating behavior (Results), one potential explanation for our findings is that selection on remating behavior is insufficiently strong or consistent to elicit a substantial evolutionary response. That is, if females are only infrequently exposed to the consequences of completed heterospecific matings, then selection on traits that mitigate these consequences could be relatively weak. In our study, only ~14% of *D. pseudoobscura* females mated with *D. persimilis* males within 3 hours of enforced co-housing, and *D.pseudoobscura* females do not produce fewer progeny when mating with heterospecific males (see methods, and Lorch and Servedio 2005). In comparison, in Matute’s (2010) study that detected enhanced remating in sympatric *D. yakuba* females, ~30% of *D. yakuba* females mated with a *D. santomea* male within a 1 hour observation period (Matute 2010, Table S4) and females produce fewer progeny in heterospecific crosses, potentially contributing to the different outcomes of that study and our data here. This relatively high first mating rate between *D. yakuba* females and *D. santomea* males should impose stronger selection on sympatric *D. yakuba* to evolve remating habits that reduce the negative effects of heterospecific matings. Alternatively, because female receptivity is also known to be influenced by the number of sperm in storage (the ‘sperm effect’; Manning 1962, 1967), *D. yakuba* sympatric females might remate more rapidly because they experience more acute sperm depletion following heterospecific matings (as inferred in Matute 2010), rather than the because of past reinforcing selection for higher remating in response to suboptimal (interspecies) matings. It is difficult to disentangle these two hypotheses without information on remating rates with conspecific males (remating in *D. yakuba* was examined only after heterospecific matings).

Alternatively, other forces might be more critical in shaping *D. pseudoobscura* remating behavior than exposure to heterospecifics. In particular, remating behaviors are determined by complex interactions between males and females, some of which might act in ways counter to reinforcing selection imposed by exposure to heterospecifics. There is substantial evidence for sexually antagonistic coevolution acting on remating traits (Parker and Partridge 1998, Arnqvist and Rowe 2005, Crudgington et al. 2005); in these cases, individuals are expected to be well equipped to respond to antagonistic measures employed by others from their own population, but potentially poorly defended against individuals with whom they have not co-evolved.

Intriguingly, our observations of remating behavior are consistent with these outcomes of local co-evolution due to sexual antagonism. We found that females mated previously to male conspecifics of their own population remated significantly more quickly and/or more frequently than females previously mated with conspecific males from a different population or with heterospecifics; remating was least frequent after mating with heterospecific males. These observations suggest that increased sexual familiarity results in females better able to combat male post-copulatory manipulation (‘molecular coercion’; Parker and Patridge 1998) via the seminal fluid in ejaculate. A similar pattern has been previously observed in Bean Weevils, in which matings involving increasingly more distantly related first males resulted in increasingly reduced rates of female remating; first matings with heterospecific males elicited the greatest post-copulatory egg production and the lowest re-mating rate (Fricke et al. 2006). In both cases, females appear to be more able to resist suppression of remating by local males in comparison to unfamiliar males.

Nonetheless, it should be noted that while our data are consistent with one scenario of sexual conflict, patterns of remating that result from sexual conflict dynamics can be complex, and are not necessarily consistently predictable from population or species crosses (Tregenza et al. 2006). In particular, previous work indicates that the outcome of interactions with less familiar males is dependent on which sex is “ahead” in any particular instance (Long et al. 2006, Tregenza et al. 2006). Our findings are consistent with females being “behind” relative to foreign males, but “ahead” of local males, however alternative patterns can be detected in these kinds of comparisons. For example, Long et al. 2006 performed crosses between six sister laboratory populations that had been isolated for 600+ generations and found within-population variation among females in whether they performed better or worse following mating with foreign males compared to local males; they concluded this was due to segregating variation for whether the female was ‘ahead’ or ‘behind’ the specific male genotype to which she was mated. These and other studies indicate that potentially more complex patterns can equally be consistent with conflict scenarios, compared to the one we infer from our observations.

Other interpretations of our finding are also possible. For example, the patterns we observed could be due to cryptic female choice for foreign or rare male sperm. This would be especially curious in heterospecific matings, as hybrid inviability makes it strongly disadvantageous for females to preferentially choose sperm from heterospecific males, and for this reason we favor the sexual conflict interpretation. Regardless, our data are clearly inconsistent with reinforcement shaping responses in this trait.

In addition to examining remating traits, our experimental design allowed us to reassess evidence of reinforcement in first matings involving these populations. As with a parallel larger study with many of the same isofemale lines (Castillo and Moyle 2016), we found no evidence for reinforcement in first mating between our populations. Discrimination against heterospecific males was not stronger in historically sympatric females, the most straightforward expectation of a response to reinforcing selection. This is curious, as previous studies have detected significantly stronger sexual isolation in sympatric *D. pseudoobscura* females (Noor 1995; Noor and Ortiz-Barrientos 2006). At least two factors could potentially contribute to our observed differences. First, sympatric populations might be polymorphic for high discrimination alleles (as suggested in Barnwell et al. 2008), and we happened to use lines that discriminate differently compared to previous studies. Second, Anderson and Kim (2005, 2006) have argued that gene flow among *D. pseudoobscura* populations has contributed to homogenizing mating discrimination traits between allopatric and sympatric sites. Interestingly, the range of mean heterospecific mating rates for sympatric isofemale lines is broad in both our analysis and in Noor’s (1995) study (range=0.22-0.52, 0.16-0.37, respectively). In addition, the sympatric lines in our study have somewhat higher heterospecific mating rates (mean=0.346) compared to Noor’s (1995) sympatric lines (mean = 0.252), whereas our allopatric lines mated with heterospecifics at considerably lower rates than Noor’s (mean=0.319 versus 0.45 in Noor 1995). These differences suggest indirect evidence that genetic polymorphism within sympatric populations and gene flow/homogenization between *D. pseudoobscura* populations might both contribute to differences between our findings and those in Noor (1995).

Regardless of these observations for first matings, our primary analysis of remating suggests that factors such as local sexual coevolution could act counter to reinforcing selection. Even when advantageous for females to manipulate the genetic identity of offspring via remating—such as following matings with heterospecific males—our results suggest that behavioral manipulation of females by male seminal proteins could supersede this response. It has been broadly recognized that sexually antagonistic coevolution and reproductive character displacement can interfere with each other, producing sub-optimal outcomes for one or both processes. Interestingly, this potential tension between intraspecific and interspecific sexual interactions is more often described in terms of reproductive character displacement hampering optimal outcomes of intraspecific sexual selection, rather than the reverse (Ortiz-Barrientos et al. 2009, Pfennig and Pfennig 2012). Here we infer that intraspecific sexual dynamics might instead overwhelm the action of reinforcing selection, producing complex outcomes for remating behaviors within and between species.

## Acknowledgements

We would like to thank M. Noor, A. Hish, and N. Phadnis for providing strains used in this experiment, and Donn Castillo for help with collecting strains. Collections were completed with assistance from IU Biology Department travel awards to DMC. Research was supported by Indiana University Dept. of Biology funding to LCM, DMC and JSD.

